# Chaperone-driven entropic separation of amyloid nanofilament bundles

**DOI:** 10.1101/2023.05.24.542046

**Authors:** Jose M. G. Vilar, J. Miguel Rubi, Leonor Saiz

**Affiliations:** Biofisika Institute (CSIC, UPV/EHU) and Department of Biochemistry and Molecular Biology, University of the Basque Country (UPV/EHU), P.O. Box 644, 48080 Bilbao, Spain; IKERBASQUE, Basque Foundation for Science, 48011 Bilbao, Spain; Departament de Fisica de la Materia Condensada, Universitat de Barcelona, Marti i Franques 1, 08028, Barcelona, Spain; Department of Biomedical Engineering, University of California, 451 E. Health Sciences Drive, Davis, CA 95616, USA

## Abstract

The disassembly of misfolded protein aggregates is a requirement for the proper functioning of cells. It has implications in multiple neuropathologies, such as Alzheimer’s and Parkinson’s diseases. The active unbundling of fibrillar aggregates has recently been identified as a key, rate-limiting step in the disassembly process. Yet, the nature of the underlying molecular mechanism remains an outstanding question. Here, we develop a coarse-grained computational approach from the atomistic structural information and show that the interactions of molecules tethered to fibrils lead to entropic forces consistent with the unbundling process observed in amyloid α-synuclein disaggregation by Hsp70. We uncover two main types of entropic effects, categorized as intra- and inter-protofilament, which are differentially affected by the system parameters and conditions. Our results show that only highly efficient chaperone systems can overcome the free energy cost of the lateral association between two protofilaments. Through the analysis of cryo-electron tomography and high-speed atomic force microscopy data, we find that co-chaperone networks and ATP hydrolysis are needed to achieve the conditions for highly efficient entropic force generation. We highlight the implications of these results for the design of targeted therapies for the underlying neuropathologies.

## Introduction

Proteins in the cellular environment are susceptible to losing their functional structure, exposing buried regions that can interact with those of other proteins and promote aggregation (1). The resulting misfolded aggregates can be amorphous (2), fibrillar (3), or a combination of both (4). They are frequently toxic, especially short fibrillar ones, leading to the disruption of normal cellular functions (5). To counteract these effects, cells have evolved specialized molecular systems that target aggregates to either degrade their components or refold them back into functional structures (6, 7). Failure to cope with aggregation has multiple pathological implications, potentially resulting in Alzheimer’s disease and synucleinopathies such as Parkinson’s disease, multiple system atrophy, and dementia with Lewy bodies (8).

There are two major types of macromolecular mechanisms engaged in the aggregate disassembly process: those that use energy from ATP hydrolysis to pull out the misfolded proteins threading through a pore (6) and those that rely on entropic effects to extract the constituent chains (7). Energetic mechanisms have been the subject of intense research. The molecular engines involved consist of ring-like oligomers that use ATP-driven coordinated conformational changes to thread the polypeptide chain from the aggregate through the central ring pore. This action effectively disrupts the interactions between the proteins, allowing them to be refolded. Single-molecule analyses have shown the pulling forces exerted by these chaperone complexes to reach values as high as 20 pN (9) and 50 pN (10).

The characterization of entropic effects has proved to be more challenging. The core process is the binding of the chaperone complex to a disordered region of the protein. In amorphous aggregates, this event prevents that disordered region from binding to the aggregate. Additional binding events to subsequently exposed regions by random thermal motion prevent larger portions of the protein from rebinding the aggregate, effectively exerting a pulling force. In fibrillar structures, chaperones bound to the disordered regions can collide with the amyloid structure, which would generate a net-pulling force away from the surface (11). Therefore, it was widely believed that misfolded proteins were extracted analogously one by one from the ends of the fibril or the surface of the aggregate through this type of entropic pulling (12). This behavior is indeed consistent with the observed exponential-like kinetics of the overall aggregated mass decay triggered by chaperone-driven entropic effects for compact amorphous aggregates, but it is not so for fibrillar structures (13).

Recent single-aggregate experiments have shown that for short fibrillar structures, of the order of 100 nm in length, disassembly occurs as an all-or-none phenomenon (14). Typically, the fibrillar structure consisting of a two-protofilament bundle remains in an apparently metastable state, with characteristic time scales of hours, before suddenly disaggregating completely, within the scale of about a minute (14-16). High-speed atomic force microscopy shows that the sudden disaggregation process progresses along the concomitant sequential unbundling of the two fibers (14). For longer fibrillar structures, disassembly occurs stepwise with all-or-none phenomena occurring sequentially in segments ∼100-nm long on average (15). The net effect in a population of aggregates is the progressive shortening of the average length of the fibrils as if it were a continuous removal of aggregate units from the fibril ends (11, 12).

All this experimental evidence indicates that forces generated by molecular chaperones can potentially be responsible for the cooperative unbundling of the constituent protofilaments of the fibrillar structure. Here, we show that force in this process is generated through a novel entropic mechanism, markedly different from the entropic forces that have been described so far in disaggregation (11, 17, 18) and in other cellular processes, such as clathrin-coat disassembly (19) and protein translocation (20, 21). To characterize the underlying mechanism, we have developed a coarse-grained approach that closely mimics the essential elements of the atomic-level molecular description. We show that, indeed, the interactions of molecules tethered to fibrils lead to effective repulsive entropic forces between protofilaments consistent with the observed unbundling process. As explicit system, we consider α-synuclein fiber-Hsp70 chaperone complexes and we compare Brownian dynamics simulation analyses with cryo-electron tomography, quantitative atomic force microscopy, and high-speed atomic force microscopy. Our results reveal two dominant entropic effects. One of them appears for short inter-protofilament distances from the cooperative interactions between tethered chaperones. The other one dominates for intermediate distances, is non-cooperative, and results from the interactions between chaperones and protofilaments. Both types of effects, but especially cooperative ones, need to be highly efficient to overcome the association forces between protofilaments. Comparing with the available experimental data, we show that co-chaperone networks and ATP hydrolysis are needed to achieve the conditions for high efficiency.

## Results

### From molecular components to coarse-grained structures

Coarse-grained descriptions are frequently used for capturing the overall behavior of biomolecular systems over longer and larger scales than what would be possible with atomistic models (22, 23). The level of detail ranges from multiple groups of atoms within a protein (24) to a single group of atoms with the appropriate geometry for the whole protein in complex biomolecular systems (25). Here, in setting the core elements of the coarse-grained description, it is crucial to realize that entropic forces are determined by the underlying system geometry and the thermal energy, which sets their strength (26).

The starting point to obtain a geometric description of the system is the available structural data. Explicitly, the fibrillar system to be disassembled consists of two bundled protofilaments. Each protofilament has a beta-sheet core of ∼5 nm in diameter, consisting of 0.5 nm thick stacked α-synuclein molecules with protruding intrinsically disordered C- and N-terminal regions (27) (Fig. 1A). In general, although there is structural polymorphism, the geometric aspects remain largely similar (28). We focus on the binding of 70-kDa heat shock proteins (Hsp70s), which are ubiquitous in protein folding and remodeling processes (7). Members of the Hsp70 family, such as Hsc70 and DnaK, share common structural motifs, including two compact domains, the substrate-binding domain (SBD) and the nucleotide-binding domain (NBD), tethered to each other through a flexible ∼20-residue long linker (29) (Fig. 1A). The SBD binds the substrate, in this case, any of the α-synuclein C- and N-terminal regions. The binding occurs with high affinity when the NBD is in the ADP-bound state. Since Hsc70 binds one every other α-syn monomer at saturating concentrations, there is a maximum of one tethered molecule per nm of protofilament (11). Therefore, the core structural unit is two α-synuclein molecules and a chaperone, which is repeated along the fiber alternating the protofilament the chaperone binds to.

**Figure 1.**
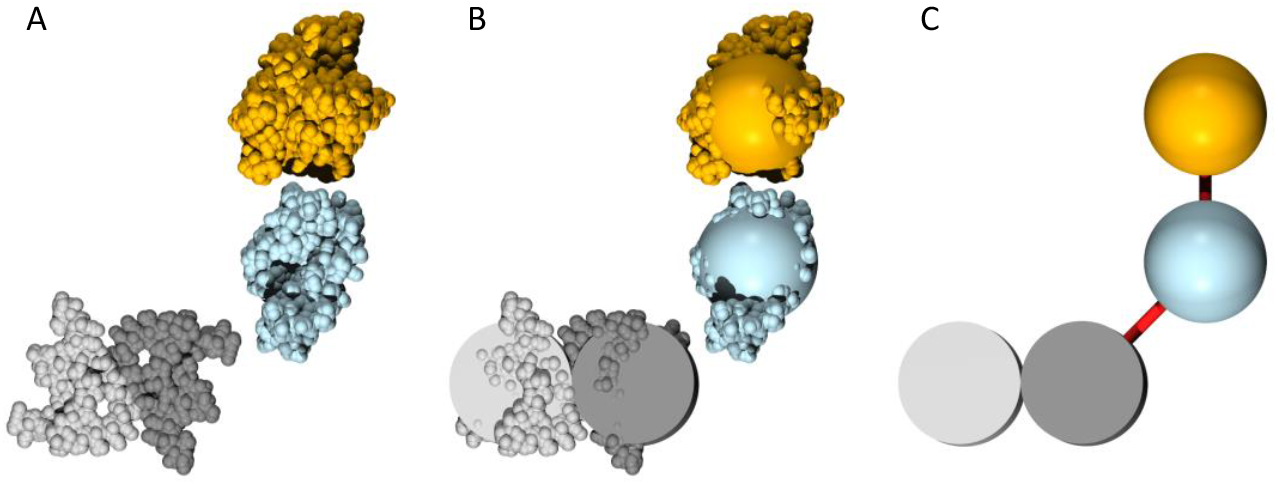
From structure to the coarse-grained representation of the α-synuclein-Hsp70 system. (A) The atomic structure of a cross-section of the structured region of a fiber is shown in gray (PDBID 6CU7) (27). Dark and light gray tones differentiate two α-synuclein molecules (residues 38-97) according to their protofilament. The structures of the NBD (residues 1-370) and SBD (residues 390-600) of Hsp70 (PDBID 2KH0) (29) are shown in orange and blue, respectively. The blue SBD can interact with the dark gray α-synuclein molecule through the intrinsically disordered regions (not shown) located at the α-synuclein tails (residues 1-37 and 98-140). (B) The coarse-grained representation is shown as geometric figures superimposed with the atomic structures. (C) The coarse-grained model used in the simulations includes two linkers (red lines). Hsp70 is represented as two 2.5 nm-radius spheres tethered to each other up to a distance of 6 nm from center to center. The SBD is tethered to one of the 2.5 nm-radius cylinders that represents an α-synuclein molecule in one of the protofilaments of the fibril up to 7 nm from center to center.

The simplest geometry that can capture the fundamental nature of the molecular system consists of each protein represented as two tethered spheres and each protofilament, as a long cylinder (Fig. 1B). The scales of these components are key parameters. Therefore, consistently with the structural information, we model the fiber as two 5 nm-diameter parallel hard-core cylinders. The SBD, modeled as a 5 nm hard-core sphere, is restricted to move within 2.0 nm of the surface of the cylinder it is attached to, or equivalently, within 5 and 7 nm from the protofilament axis to its center position. Similarly, the NBD is modeled as a 5 nm hard-core sphere restricted to move within 1 nm of the SBD it is tethered to, which corresponds to a range within 5 and 6 nm center-to-center distance (Fig. 1C).

### Optimal disaggregation

We performed Brownian dynamics simulations of the coarse-grained system to characterize the forces exerted by the random thermal motion of the chaperones attached to the protofilaments. We considered first the observed optimal experimental conditions and subsequently suboptimal conditions and modified elements.

Optimal disaggregation involves saturating concentrations of chaperones, with the observed stoichiometry of one Hsc70 molecule bound every two α-synuclein molecules. Therefore, we attached the chaperones to the protofilaments spaced 1 nm from each other with an offset of 0.5 nm between protofilaments. We considered 200 chaperones, which resulted in a 100 nm coverage of the fiber, and started with the closest distance possible between protofilaments, namely, a 5 nm axis-to-axis distance. Experimentally, it has been observed that 100 nm fibers at saturating conditions lead to all-or-none disaggregation for 90% of the fibrils in about three hours (14). As initial conditions, we placed the chaperones attached to each protofilament, fully stretched with a random orientation around the external half-side of the protofilament, and perpendicular to the protofilament axis. In this way, chaperones attached to one protofilament are separated from chaperones attached to the other protofilament and they occupy the maximum volume available to them (Fig. 2A,E).

**Figure 2.**
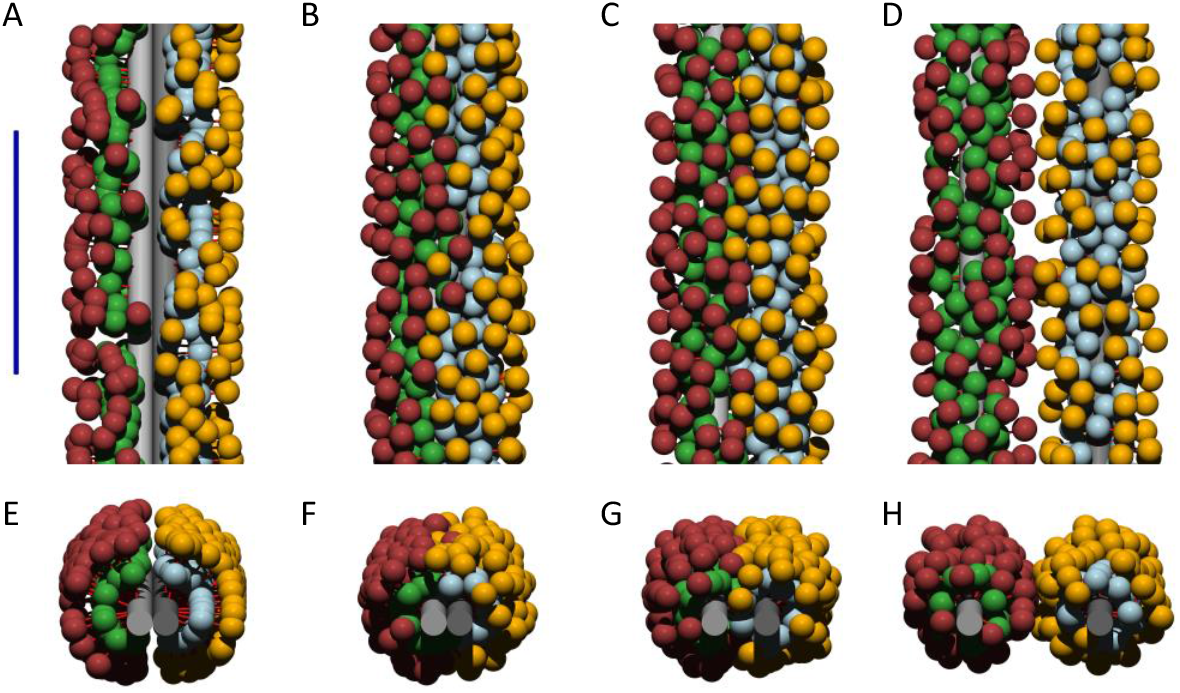
Restrictions in chaperone degrees of freedom are lifted progressively as the protofilaments move further apart from each other. Representative state snapshots of the coarse-grained Brownian system are shown for protofilament axes distances of (A,E) 5 nm with the initial conditions; (B,F) 5 nm after initial conditions relaxation, which maintains chaperone separation; (C,G) 10 nm, which allows for SBDs and NBDs to be placed between protofilaments; and (D,H) 25 nm, which allows for almost free motion of chaperones around the protofilaments. SBDs and NBDs bound to the left (right) protofilament are colored green (blue) and red (orange), respectively. The top and bottom panels show top and front views of the amyloid bundles, respectively. The blue scale bar in panel A corresponds to 50 nm.

After equilibration, steric clashes present in the initial conditions are resolved, with the chaperones remaining tightly packed and spatially separated according to the protofilament they attached to (Fig. 2B,F). Therefore, the stoichiometry observed at saturation corresponds essentially to the maximum density of chaperones that can be achieved given the geometric constraints. The system can maintain the separation of the chaperones bound to each protofilament, restricting their motion through collisions with either the opposite protofilament or the chaperones attached to it. This restriction is lifted progressively as the protofilaments move further apart from each other (Fig 2C,G and 2D,H), resulting in states with increasingly higher entropy. Under these conditions, protofilaments are expected to experience an effective repulsive force toward higher entropy states.

Indeed, the forces exerted by chaperones on each protofilament when they are next to each other act in opposite directions (Fig. 3A). The net effect is an effective repulsion force of ∼3 pN. A key property is how far apart this force can be sustained to counteract the attractive interactions that keep the protofilaments in a bundle (30, 31). Our results show that the force remains virtually unchanged from 5 to 10 nm, with an average value of 2.9 pN. At 10 nm, there is a sharp decrease in the repulsive force, which remains at 1.3 pN on average up to a separation of 15 nm and monotonously decays to zero further apart. This force profile results in a free energy change of −23.8 pN nm for the overall unbundling process (Fig. 3B).

**Figure 3.**
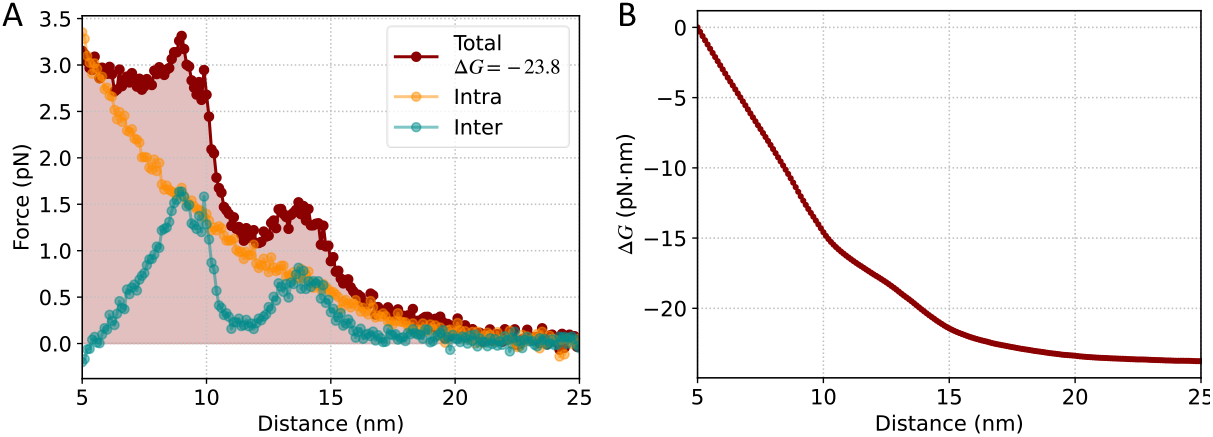
Forces exerted by chaperones on each protofilament act in opposite directions. (A) The total entropic force exerted as a function of the separation between protofilaments (red) and its decomposition in intra-protofilament (orange) and inter-protofilament (dark cyan) components are shown for a system with 200 chaperones, 2 chaperone/nm density, 6 nm SBD-NBD tether, and 7 nm protofilament-SBD tether. The free energy change for the overall unbundling process is shown in the legend in units of pN nm and (B) as a function of the separation between protofilaments.

### Two types of effects dictate disaggregation progression

To elucidate the potential origin of the non-monotonous force profile, we analyzed the forces acting on a protofilament by breaking them down into two categories: those coming from chaperones tethered to the same protofilament, which include both pulling and collision effects, and those coming from chaperones tethered to the opposite protofilament, which consist of just collisions (Fig. 3A). The intra-protofilament forces from the chaperones tethered to the same protofilament decrease monotonically as a function of the distance whereas the inter-protofilament forces from the chaperones tethered to the opposite protofilament are responsible for the sharp transitions. They show two maxima around protofilament separations of 10 and ∼15 nm. The maximum at 10 nm is coincidental with one or two of the chaperone domains placed between the protofilaments. Two domains require the molecule to be parallel to the protofilament axes. As the two domains become perpendicular to the axes, they would match the 15 nm separation. Remarkably, the inter-protofilament effects, although small in absolute value, are negative when the fibrils are next to each other. The negative values arise from chaperones being able to extend over the opposite protofilament they are attached to, which results in collisions on the external half-side of the opposite protofilament. Therefore, intra-protofilament effects are needed for the initial unbundling and inter-protofilament effects, for the subsequent progression.

### System size effects

Cryo-electron tomography of fiber-chaperone complexes has shown predominantly densely clustered chaperones over extended stretches of 30-100 nm on fibers incubated with non-saturating concentrations of Hsc70. Therefore, system size may be relevant for the disassembly process. In addition to 200 chaperones bound to a 100 nm stretch, we considered 100 and 20 chaperones bound to a 50 and 10 nm stretch, respectively (Figs. 4A and 4B). We also investigated larger systems at the same density by considering 1,000 chaperones bound to a 500 nm stretch. The results show that, at short distances, the entropic force per chaperone increases with the system size (Figs. 3 and 5). However, the differences are minimal for relatively large systems (50, 100, and 500 nm). A small system, such as 20 chaperones bound to a 10 nm stretch, in contrast, is very inefficient at short distances: not only do intra-protofilament effects decrease but also inter-protofilament effects become more negative (Fig. 5C). The free energy changes are −10.9 pN nm per chaperone for the 10 nm system compared to −20.1, −23.8, and −27.5 pN nm for the 50, 100, and 500 nm systems, respectively.

**Figure 4.**
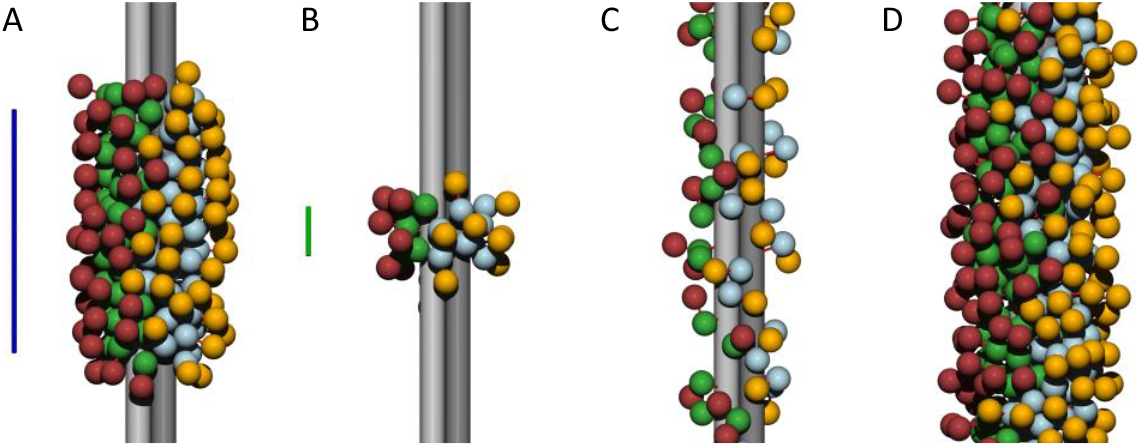
From optimal to inefficient systems. Representative state snapshots of the coarse-grained Brownian system at protofilament axis distances of 5 nm after initial conditions relaxation for systems as in Fig. 2B except for (A) 100 chaperones; (B) 20 chaperones; (C) 0.4 chaperone/nm density; and (D) 7 nm SBD-NBD and 9 nm protofilament-SBD tethers. The restrictions in the chaperone degrees of freedom decrease as the system size decreases, the chaperone density decreases, or the tether length increases. The blue and green scale bar in panels A and B corresponds to 50 and 10 nm, respectively.

**Figure 5.**
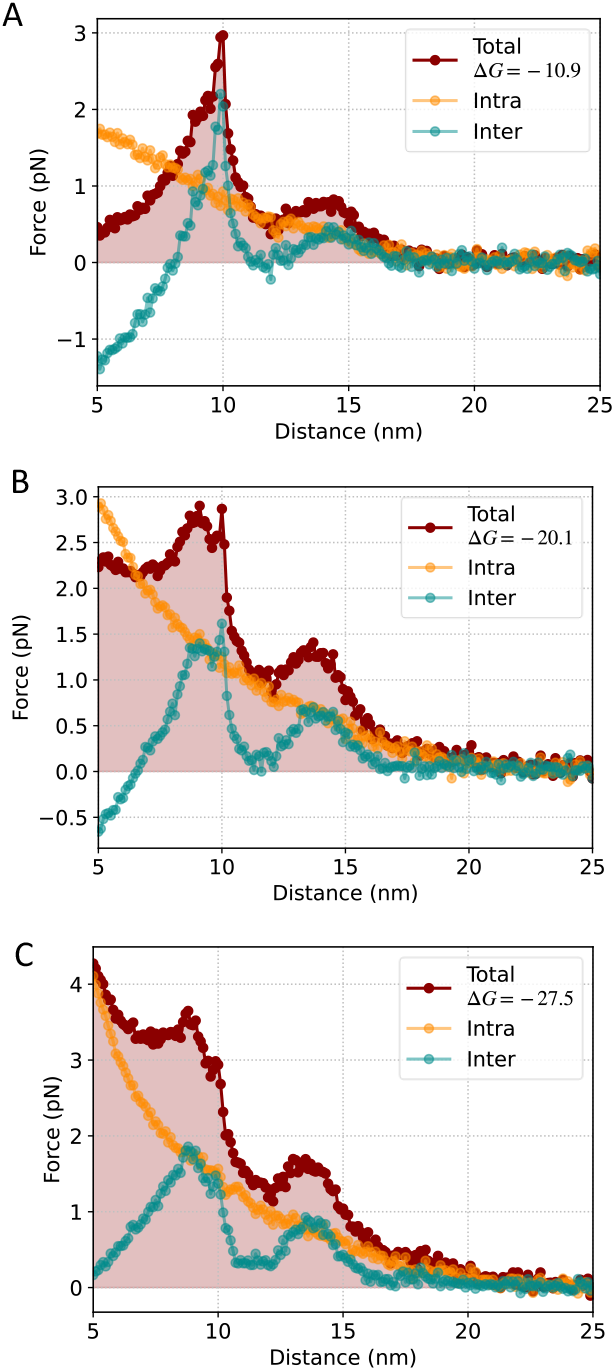
The overall effect of entropic forces decreases as the system size decreases. Entropic forces and total free energy changes per chaperone are shown under the same conditions as in Fig. 3A except for system sizes of (A) 20 chaperones; (B) 100 chaperones; and (C) 1000 chaperones. Not only do intra-protofilament forces decrease but also inter-protofilament forces become more negative at short distances as the system size decreases.

### Chaperone density effects

Collective effects, namely, how the force exerted per chaperone depends on the density of chaperones bound to the protofilaments, are expected to play a fundamental role. To investigate this aspect, we performed simulations at low densities, considering explicitly one chaperone every 2.5 nm on the fibrillar bundle, alternating between protofilaments (Fig 4C). Under these conditions, the intra-protofilament effects disappear almost completely whereas the inter-protofilament effects are maintained at high absolute values, switching from negative to positive as the separation between protofilaments increases (Fig. 6). These results reveal that high density of chaperones over stretches of 30-100 nm on the fiber, as observed experimentally, is strictly needed for chaperones to be able to exert a repulsive force when the protofilaments are next to each other.

**Figure 6.**
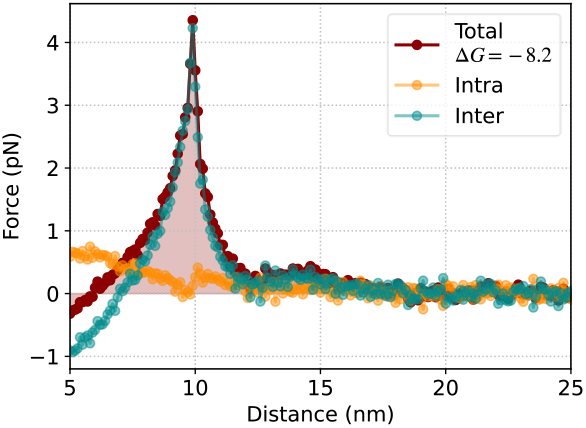
Intra-protofilament entropic forces become negligible at low chaperone density. Entropic forces and total free energy changes per chaperone are shown under the same conditions as in Fig. 3A except for a density of 0.4 chaperones/nm (1 chaperone every 2.5 nm). Only collisions of the chaperones with the opposite protofilament are relevant. The inter-protofilament force achieves higher values at the resonance distance of 10 nm than in Fig. 3A because the two domains of an Hsp70 molecule can enter the space between the protofilaments oriented parallel to their axes with lower constraints in their orientation. At high densities, the perpendicular orientation dominates.

There are notable similarities in the behavior of small and low-density systems (Figs. 5B and 6). It is so because small systems cannot achieve a high effective chaperone density due to boundary effects. Densely packed chaperones are restricted in their movement by collisions with neighboring chaperones and tend to stay close to their attachment point. However, at low densities or at the edges of a densely clustered system, chaperones can extend their linkers along the protofilament. For example, in a system with 20 chaperones attached to a 10 nm stretch of protofilament, the chaperones extend over ∼20 nm (Fig 4A). This inefficiency in small systems or low densities provides a rationale for the observed clustering of chaperones over ∼30-100 nm stretches.

### Alternative system parameters

A key determinant of the system behavior is the effective volume available to the different degrees of freedom. In addition to lowering the density along the fiber, we controlled this aspect by changing the length of the tethers. Since the system is densely packed, it was only feasible to increase their length (Fig 4D). By increasing the length of the protofilament-SBD and SBD-NBD tethers by 2 and 1 nm, respectively, the forces exerted are reduced substantially (Fig. 7). These changes are reminiscent of those for small systems and low density (Figs. 5A and 6) but the origin of this effect is different. In particular, the presence of a longer tether, and concurrently less stringent packing, allows the chaperone cloud of each protofilament to be mixed with each other to a larger extent. Therefore, the motion is not as restricted as with shorter tethers. In terms of the chaperone distribution, it manifests as an interface between the two clouds of chaperones rougher for long tethers (Fig. 4D) than for short ones (Fig. 2B).

**Figure 7.**
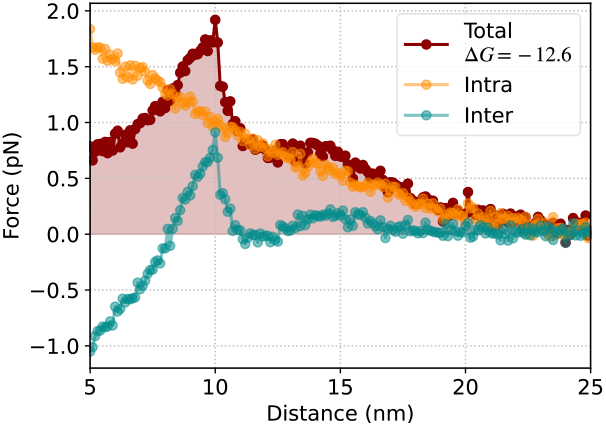
Changes in key system parameters strongly affect entropic forces. Entropic forces and total free energy changes per chaperone are shown under the same conditions as in Fig. 3A except for longer tethers (7 nm SBD-NBD and 9 nm protofilament-SBD tethers). Longer tethers imply looser packing and smaller entropic force. Inter-protofilament forces can become clearly negative at short distances.

### Quantitative experimental validation

The overall dimensions of the coarse-grained geometry obtained through Brownian dynamics simulations (Figs. 2B and 2C) are also consistent with those of the system as observed through cryo-electron tomography of fiber-chaperone complexes (Fig. 3D of Benton et al. (15)), which shows a ∼35 nm wide chaperone-decorated bundle. The similarities of Figs 2B, and 4A with the experimental data are remarkable. The experimental data also shows that the chaperones are positioned close to the protofilaments.

The force profile as a function of the separation between protofilaments is consistent with the unbundling process observed through high-speed atomic force microscopy. Generally, there is a fast depolymerization as the protofilaments get unbundled (14). However, there are cases in which the two protofilaments remain parallel during the unbundling process long enough to be observed. For instance, Fig. 3E of Franco et al. (14) shows two 100 nm long protofilaments that remain parallel to each other at a distance of 20 nm for over 20 s before depolymerizing. The value of 20 nm is consistent with the distance at which entropic forces become negligible (Fig. 3).

Quantitative atomic force microscopy has been used to characterize the energetics of α-synuclein fiber formation (31). The results show a free energy change of the lateral association between two protofilaments in a fibril to be between −18.3 and −12.3 pN nm per pair of α-synuclein molecules. We obtain a free energy change of −23.8 pN nm per chaperone, which at saturation corresponds to a pair of α-synuclein molecules, for the unbundling process under optimal conditions (200 chaperones over 100 nm). Therefore, our results show that indeed entropic forces have enough strength to drive the unbundling process but only close to the optimal conditions. Suboptimal chaperone arrangements, as for instance small systems, low density, or longer tethers would not generate enough force for the unbundling to occur. Chaperone density is particularly relevant because it contributes to both the force generated per chaperone and the number of chaperones per unit of length. To overcome the association of two protofilaments, both contributions are needed.

## Discussion

The unbundling of fibrillar structures has been shown to be a key pathway in the disassembly of misfolded protein aggregates (14). A major question that remained was the mechanism behind this process. We have shown that molecules attached to the protofilaments of the bundle with the geometric constraints imposed by the atomic structural information lead to effective repulsive forces between the protofilaments strong enough for the unbundling to progress completely. The resulting forces are about 3 pN per chaperone when the protofilaments are next to each other and result in free energy changes of −23.8 pN nm for the overall unbundling process of the prototypical experimental system. The forces originate as a net statistical effect from the random thermal movements of the chaperones attached to one protofilament toward higher entropy states in which their motion is not impaired by the other protofilament and the chaperones attached to it.

The collective behavior of the system is far from trivial, even in its most fundamental aspects. Overall, our results show a highly efficient system, but only under specific conditions. The requirements for efficiency include chaperones present at sufficiently large numbers, packed at high densities, and restricted to move close to the protofilament they are attached to. All these requirements have been observed simultaneously in functioning systems (15).

High density of chaperones is needed when the protofilaments are in close proximity because the repulsive component of the entropic force is dominated by the collisions of the chaperones with each other. Therefore, it is a highly cooperative effect at short distances. Indeed, fibers incubated with non-saturating concentrations of Hsc70, DNAJB1, Apg2, and ATP show segments of densely clustered chaperones stretching in length typically from 30 to 100 nm. Namely, the biological system has been designed to either have long regions with either a high density or no chaperone at all.

The functional involvement of co-chaperones, such as DNAJB1 and Apg2, is necessary for the effective operation of the system. Specifically, in the presence of ATP, DNAJB1 helps the binding of Hsc70 to α-synuclein fibers, and Apg2 triggers further chaperone binding and clustering from the end of the fiber (15). Our results demonstrate that without this high-density clustering, the repulsive force per chaperone would be minimal at short distances. The presence of chaperones at sufficiently large numbers is a requirement because small systems cannot achieve high densities. Establishing all these optimal conditions is energetically costly in terms of ATP hydrolysis. The presence of ATP, however, does not seem to impart an enthalpic component to the forces generated by Hsp70 (32).

At larger inter-protofilament distances, there is a major contribution to the unbundling force from the interactions of the chaperones with the opposite protofilament. We identified a spatial resonant-like phenomenon at the effective size of the chaperone domains. This effect is due to the interactions of one or two domains with the two protofilaments as they localize between the protofilaments. At the first resonant distance, the magnitude of the force per chaperone is about 3 pN. Interactions with the opposite protofilament, however, can also have a negative impact on disaggregation if chaperones can collide with its external half-side. Indeed, this situation can happen when the motion of the chaperones is not restricted to be close to the protofilament they are attached to, as for instance when tethering lengths increase.

The characteristic forces up to the first resonant distance are comparable to the maximum forces per protomer achievable by the chaperone complexes that hydrolyze ATP to generate mechanical force, which in the case of the hexameric ClpX and ClpB chaperones can reach values of 20 and 50 pN, respectively (9, 10). Overall, entropic forces lead to free energy changes of −23.8 pN nm per chaperone which can overcome the −18.3 and −12.3 pN nm per pair of α-synuclein molecules obtained for the lateral association between two protofilaments (31).

The existence of different mechanistic regimes, namely short-range cooperative and intermediate-range resonant, and the requirements for functionality make the understanding of the disassembly process an essential step in the development of targeted therapies for the underlying neuropathologies. Particularly relevant are the design principles we have uncovered, including the dependence of the different contributions on the chaperone density on the fibril and the impact of the length of the tethers, embodied by the interdomain linker and attachment points at the protruding intrinsically disordered C- and N-terminal regions of α-synuclein, which provide essential information for the rational development of targeted approaches.

## Methods

### Dynamic equations

We performed Brownian dynamics simulations (22) of the system’s coarse-grained description to characterize the random thermal motion of the chaperones attached to the protofilaments. In this framework, the displacement Δ**r**_*i*_ of a particle *i* in a time interval Δ*t* is given by

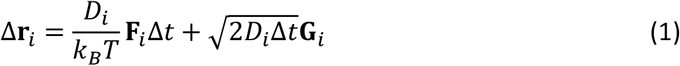

where the subscript *i* refers to the particle index, *D*_*i*_ is the translational diffusion coefficient of the particle, *F*_*i*_ is the force acting on the particle, *k*_*B*_ is the Boltzmann constant, *T* is the absolute temperature, and **G**_*i*_ is a normally distributed random vector. The Stokes-Einstein relation, *D*_*i*_ = *k*_*B*_*T*/6*πηR*_*i*_, leads to values of *D*_*i*_ = 0.098 nm^2^ ns^−1^ for *R*_*i*_ = 2.5 nm spheres in water at 25°C (33). Note that, at 298 K, the thermal energy is *k*_*B*_*T* = 4.114 pN nm.

### Coarse-grained forces

The force acting on particle *i* comprises hard-core and tethering interactions. Hard-core interactions between protein domains *i* and *j* have radial symmetry and are described mathematically through

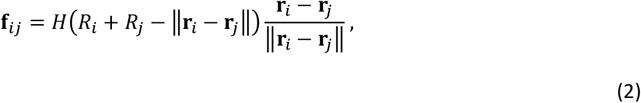

where *H*(*r*) is zero for *r* < 0 and infinity otherwise. Hard-core interactions between protein domains and protofilaments are perpendicular to the protofilament axis, which is captured through the expression

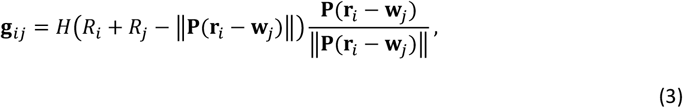

where *i* and *j* refer to the protein domain and protofilament, respectively, 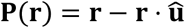 indicates the projection of the vector **r** in the plane perpendicular to the direction of the fiber, described by the unit vector 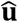. Here, **w**_*j*_ is any position along the axis of the protofilament.

Tethering restricts how far apart particles can be from each other. There are tethering between the two domains of a protein and between the SBD particles and the protofilament they are bound to. In both cases, tethering is described through

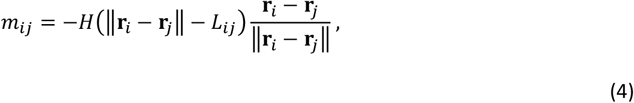

where *L*_*ij*_ represents the length of the tether between particles *i* and *j*. In the case of protofilaments, **r**_*j*_ represents the position of the chaperone attachment.

We consider only translational motion of the protein domains since the coarse-grained spherical domain and the tether have radial symmetry.

### Computational implementation

To implement the simulations, the function *H*(*r*) is approximated as

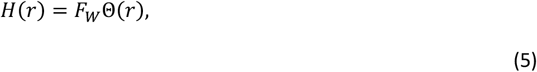

where Θ is the Heaviside unit step function and *F*_*W*_ is the force intensity. Within this framework, the average excess distance into the hard-core region or outside the tether length is Δ*r* ≃ *k*_*B*_*T*/*F*_*W*_. The value of *F*_*W*_ should be chosen large enough so that Δ*r* ≪ *R*_*i*_ + *R*_*j*_ and Δ*r* ≪ *L*_*ij*_. For instance, *F*_*W*_ = 100 pN, which leads to Δ*r* ≃ 0.04 nm, would keep the offset length below 1% of the center-to-center distances in a collision. Additionally, the value of Δ*t* should be chosen small enough to be consistent with the continuous limit. This value depends in turn on the excess distance as Δ*t* ≪ Δ*r*^2^/2*D*, which leads to Δ*t* ≪ 0.008 *ns* for *F*_*W*_ = 100 pN.

The initial randomly orientated distribution of chaperones attached to each protofilament was equilibrated over 2 μs. To facilitate equilibration, the intensity of the force was increased linearly in time during this interval from *F*_*W*_ = 1 pN to *F*_*W*_ = 100 pN. Concomitantly, the time step was decreased quadratically from Δ*t* = 0.01 ns to Δ*t* = 0.001 ns.

### Force profile characterization

The force acting on the protofilaments is computed as the average of **F**_*i*_ over time. The impulsive character of the hard core and tether approach implies that each collision or restriction would contribute a large amount over a short time. This amount becomes larger as *F*_*W*_ increases but the time it acts becomes shorter, leading to a consistent value of the average force independent of *F*_*W*_ provided that *F*_*W*_ is large enough.

To characterize the dependence of the entropic force on the separation between protofilaments, we computed the forces exerted by the chaperones for a range of distances. Explicitly, from the initial separation, we computed the average forces for 100 ns, increased the separation by 0.1 nm, let the system equilibrate for 100 ns, and iterated this process to reach a separation of 25nm between protofilaments.

To reduce the statistical fluctuations in the computed force profile to the same level in all the cases, we averaged over replicas of the system to reach a constant number of 1000 chaperones, ranging from 1 replica for the 500 nm system to 50 replicas for the 10 nm system.

## Acknowledgments

J.M.G.V. acknowledges support from Ministerio de Ciencia e Innovación under grant PID2021-128850NB-I00 (MCI/AEI/FEDER, UE).

